# Origin of eukaryotic-like Vps23 shapes an ancient functional interplay between ESCRT and ubiquitin system in Asgard archaea

**DOI:** 10.1101/2023.07.25.550441

**Authors:** Zhongyi Lu, Siyu Zhang, Yang Liu, Runyue Xia, Meng Li

**Affiliations:** Archaeal Biology Center, Institute for Advanced Study, Shenzhen University, Shenzhen 518060, China; Shenzhen Key Laboratory of Marine Microbiome Engineering, Institute for Advanced Study, Shenzhen University, Shenzhen 518060, China

**Author notes:** Address correspondence to Meng Li,. These authors contributed equally to this work (Zhongyi Lu, Siyu Zhang, Yang Liu).

**Keywords:** Asgard archaea, endomembrane system, ESCRT, ubiquitin system

## Abstract

Functional interplay between the endosomal sorting complexes required for transport (ESCRT) and ubiquitin system underlies the ubiquitin-dependent sorting pathway, a specific trait of eukaryotic endomembrane system. Yet, its evolutionary origin remains unclear. Here, we show that a novel UEV-Vps23 family protein, that contains UEV and Vps23 domains, mediates an ancient ESCRT and ubiquitin system interplay in Asgard archaea. The UEV binds ubiquitin with high-affinity, making the UEV-Vps23 a sensor for sorting ubiquitinated cargo. A steadiness box in the Vps23 domain undergoes ubiquitination through a eukaryotic-like Asgard E1, E2, and RING E3 cascade. The UEV-Vps23 can switch between autoinhibited and active forms, by which likely regulates the ESCRT and ubiquitin system interplay. Furthermore, the shared sequence and structural homology among the UEV-Vps23, eukaryotic Vps23 and archaeal CdvA, implying that these proteins share a common evolutionary origin. Together, this work presents evidence that the ESCRT and ubiquitin system interplay had arisen early in Asgard evolution, antedating emergence of endomembrane system in eukaryogenesis.

**Teaser:** The ESCRT and ubiquitin system interplay, a specific trait of eukaryotic endomembrane system, likely inherited from an Asgard archaeal ancestor.

## Introduction

Eukaryogenesis is a major puzzle in evolutionary biology, because the evolutionary origin of many eukaryotic-specific traits, especially endomembrane system, remains poorly understood. It has been thought that the endomembrane system not only leads to the intracellular compartmentalization, but also endows the intracellular membrane traffic with a ubiquitin-dependent sorting pathway (*1–3*). This pathway largely relies on a functional interplay between endosomal sorting complex required for transport (ESCRT) and ubiquitin system (*4–7*). The ESCRT is multisubunit complex that contains well-characterized subcomplexes ESCRT-I, II, III, IV, and Vps4, mediating a variety of membrane scission events in eukaryotic cells (*8*). In a canonical endocytic sorting process in *Saccharomyces cerevisiae,* for instance, the ESCRT-I subunit Vps23 captures ubiquitinated cargo proteins via a ubiquitin-conjugating enzyme variant (UEV) domain. The Vps23 then interacts with downstream Vps28 and Vps37 subunits, respectively, to complete ESCRT-I (*9*). For assembly of ESCRT-II, the subunit Vps36 of ESCRT-II firstly binds to the Vps28 using a zinc-finger insertion domain, and thereby determines the localized precision of the interacting partners Vps22 and Vps25 (*10, 11*). Finally, the ESCRT-III is recruited to the target site through the interaction of its subunit Vps20 with the Vps25, and eventually carries out membrane scission in a Vps4-dependent manner to cause ubiquitinated cargoes into invaginating buds (*12*).

Functionally, the Vps23 initiates the ESCRT and ubiquitin system interplay in most eukaryotic lineages (*10, 13*). The integral Vps23 contains the UEV domain and a Vps23 domain; the latter domain is further structurally characterized by consisting of a coiled-coil stalk region, and a helical hairpin structure steadiness box (SB) that interacts with the Vps28 (*14–17*). The SB can be modified by a cascade of ubiquitination enzymes that include a ubiquitin-activating enzyme E1, a ubiquitin-conjugating enzyme E2, and several certain Really Interesting New Gene (RING) ubiquitin-ligases E3 (*18–20*). By ubiquitin modification, excess Vps23 is degraded through the ubiquitin-proteasome pathway in eukaryotic cells (*21*). Notably, while the UEV belongs to the E2 protein superfamily that is thought to derive from an ancient bacterial ancestor, the origin and evolution of Vps23 domain remain unknown (*14*).

It has been documented that a cell division (Cdv) machinery, that includes ESCRT-III (CdvBs) and Vps4 (CdvC) homologs, is identified in some crenarchaeal orders, including *Sulfolobales*, *Desulfurococcales*, and in Thaumarchaea within TACK superphylum (*22, 23*). Since the crenarchaea lack a eukaryotic-like ubiquitin system, the Cdv machinery seems to be functionally limited to the cell division and vesicle formation (*22, 24, 25*). Moreover, the Cdv machinery utilizes a specific membrane-binding CdvA subunit, which functions instead of Vps23, to initiate assembly of CdvB and CdvC (*26, 27*). The CdvA, which has a novel coiled-coil CdvA domain, shows no significant sequence or structural similarities with the eukaryotic ESCRT-I subunits, and thus illuminates an evolutionary and functional gap between the Cdv machinery and eukaryotic ESCRT (*28*).

In recent years, Asgard archaea phylum, the closest prokaryotic relatives of eukaryotes, has been recognized to possess both the ESCRT and ubiquitin systems (*29–35*). It has been found that several Asgard genomes encode a UEV-Vps23, that is a eukaryotic Vps23 homolog, forming complexes with an Asgard ubiquitin and a Vps28 homolog (*29–34, 36*). These findings suggest that the UEV-Vps23 can functionally link the ESCRT to the ubiquitin system in Asgard archaea, leading to the hypothesis that the ubiquitin-dependent sorting pathway of eukaryotic endomembrane system was originally inherited from an Asgard ancestor. Nevertheless, molecular mechanisms, especially those of the UEV-Vps23 involved the Asgard ESCRT and ubiquitin system interplay remain obscure, which makes this hypothesis some uncertainties. Hereby, we investigate the role of the UEV-Vps23 in the Asgard interplay by bioinformatic and biochemical analyses. Combined with the phylogenetic analysis, sequence comparison and structural modeling, we also explore the evolution of the UEV-Vps23, providing evidence for its evolutionary relationships with the CdvA and Vps23.

## Results

### Ubiquitin recognition of UEV-Vps23 across Asgard lineages

As described in our previous work(*34*), a eukaryotic-like Vps23 family protein (termed UEV-Vps23, belongs to asCOG005829 and asCOG007313) that comprised of a UEV domain and a Vps23 domain (contains a stalk region and SB), has been identified in Asgard archaea. It has also been documented that this family includes a set of novel UEV-Vps28 subunits, which are featured by fusion of a Vps28 domain to the C-terminus of UEV-Vps23, in certain genomes of Heimdallarchaeia only (e. g. Hemidallarchaeota_AB_125, NCBI accession number OLS31938.1) (*36*).

To explore the evolution of Asgard UEV-Vps23, we constructed a maximum likelihood phylogenetic tree, including the UEV-Vps23 family and its eukaryotic counterparts, and a novel Asgard UEV protein group (UEV_X, e. g. NCBI accession number OLS29001.1), which contains a UEV domain, and followed by a PF05120 domain (residues 170–210, with 65.2% probability using HHpred analysis) in Gerd-, Heimdall-, and Kariarchaeia. We noted that all the identified Asgard lineages, except Loki- and Thorarchaeia, possess the UEV-Vps23, which suggests that the UEV-Vps23 likely emerged in the last Asgard common ancestor. The resulting phylogenetic tree supports the monophyly of UEV-Vps23 of each Asgard lineage with high bootstrap values, largely ruling out the possibility of horizontal gene transfer (Fig. 1a). The UEV proteins appear to be closely related to the UEV-Vps23, and hence implies that they might derive from the UEV-Vps23 family.

**Fig. 1.**
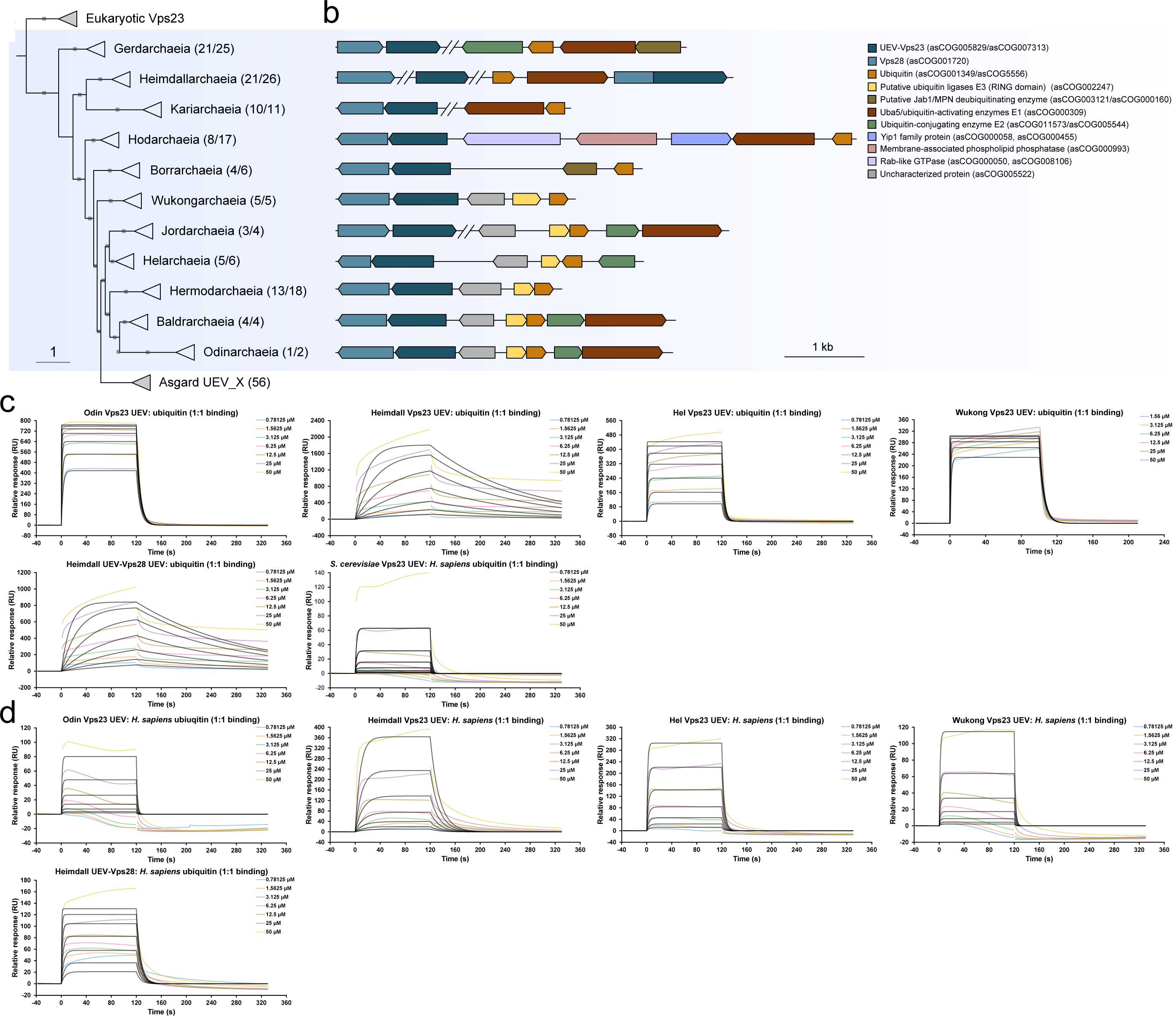
The distribution of UEV-Vps23 involved in sensing ubiquitin among Asgard lineages. (a) Phylogenetic analysis of the UEV-Vps23. The maximum likelihood tree was reconstructed by IQ-tree (settings: -m Q.yeast+F+I+G4 -bnni -B 1000), and the bootstrap values higher than 80 are shown with grey squares on tree branches. Numbers noted after the taxonomic names indicate the numbers of genomes that contain the representative gene clusters in the collapsed clades. The Heimdallarchaeia UEV-Vps28 is marked by asterisk. (b) Schematic overview of representative ESCRT and ubiquitin system component gene clusters identified in Asgard genomes. Genes from incontiguous contigs from Asgard genomes are represented with a double line at the end of the contig. The UEV-Vps28 is marked with an asterisk. (c) SPR analysis of interactions between Asgard UEV and ubiquitin. The interaction between *S. cerevisiae* UEV and *H. sapiens* ubiquitin is used as control. (d) SPR analysis of interactions between Asgard UEV and *H. sapiens* ubiquitin.

Examination of the genomic context of the UEV-Vps23 gene shows that it is part of a gene cluster that contains genes encoding a ubiquitin, a putative RING E3 ubiquitin ligase, and an uncharacterized protein (asCOG005522, e. g. NCBI accession number WEU39992.1) that is similar to the PDB: 6V1V, a vegetative insecticidal protein (residues 16–168, with 94.39% probability using HHpred analysis), in Hermod-, Odin-, Baldr-, Hel-, and Wukongarchaeia (Fig. 1b) (*34*). The vegetative insecticidal protein is identified as a secretable protein superfamily in bacteria (*37*). Borrarchaeia have a similar gene cluster organization, while its RING E3 gene is replaced by a gene encoding putative Jab1/MPN deubiquitinating enzyme (*38*). In addition to the UEV-Vps23 and ubiquitin, the gene cluster of Hodarchaeia consists of genes encoding a Yip 1 protein and a Rab-like GTPase, both of which are predicted to be involved in intracellular transport (*39, 40*).

Nonetheless, it is observed that genes encoding ubiquitin and UEV-Vps23 were not clustered in Kari-, Gerd-, Jord/LC30-, and Heimdallarchaeia, whereas the UEV-Vps28 indeed co-locates with the ubiquitin in Heimdallarchaeia. To gain further insight into the association between ESCRT and ubiquitin system, we assessed the Hermod- and Helarchaeia genomes as examples and found that genes encoding both molecular machineries are tightly clustered (Fig. S1). In eukaryotic cells, a weak affinity interaction between the UEV of Vps23 and ubiquitin has been thought to be suitable for rapid and reversible transferring the ubiquitinated cargo under multiple regulatory controls (*41*).

In order to characterize the binding capability between Asgard UEV and ubiquitin, we performed surface plasmon resonance (SPR) experiments using the Asgard ubiquitin and the UEV of UEV-Vps23 and UEV-Vps28. For a control for the Asgard interactions, we included a eukaryotic interaction between *S. cerevisiae* UEV and *Homo sapiens* ubiquitin (shares 96% of amino acid sequence similarity with *S. cerevisiae* ubiquitin). Our SPR results support the interaction between the Asgard UEV and ubiquitin, fitting to a 1: 1 binding model, with higher affinities (*K*_D_ 6.74e^-7^–7.15e^-6^ M) than that of the eukaryotic interaction (*K*_D_ 3.79e^-2^ M; Fig. 1c, Table S1). These high binding affinity of Asgard interactions imply that the UEV-Vps23 concentrates ubiquitinated cargos in a primordial mode without flexible and multiple regulations.

To test the robustness of the Asgard UEV, we determined the interaction between Asgard UEV and *H. sapiens* ubiquitin. The SPR analysis indicates that the Asgard UEV could bind *H. sapiens* ubiquitin, with the affinity being 10–1000-folds weaker than those of Asgard interactions (Fig. 1d, Table. S1). However, such SPR values for Asgard-eukaryotic interactions are still higher than those of eukaryotic interactions. These data suggest a functional compatibility of Asgard UEV architecture for sensing ubiquitin.

### Modification of UEV-Vps23 by an Asgard ubiquitination cascade

Given the presence of cluster of UEV-Vps23 and ubiquitin system components gene, we asked whether the UEV-Vps23 is a substrate of the putative ubiquitination cascade. To address this question, we identified and selected a gene operon in a Helarchaeia contig (gnl_UU_Hel_GB_B_4), which encodes complete ESCRT components and ubiquitin system that contains E1, E2, RING E3 homologues, and a pro-ubiquitin (with a Ser-Pro-Lys-Asn peptide extending beyond the conserved C-terminal di-glycine motif; Fig. 2a, Fig. S2). We expressed the E1, E2, and the mature ubiquitin (the C-terminal Ser-Pro-Lys-Asn was removed) in *Escherichia coli* BL21, and then purified and examined their functional activities in a ubiquitination reaction as previously described (*42*). The SDS-PAGE assay shows that the ubiquitination reaction that contains E1, E2, E3, ubiquitin, and full-length UEV-Vps23, resulted in the generation of potential ubiquitinated products above UEV-Vps23 band (Fig. 2b, Fig. S3a). To further confirm the C-terminal SB helical hairpin is essential for UEV-Vps23 ubiquitination ^30^, we constructed a Helarchaeia UEV-Vps23 (SB*Δ*) substrate (lacks residues 201–258), and analyzed its ubiquitination state. The SDS-PAGE and mass spectrometry assays indicate that no ubiquitinated product band of the UEV-Vps23 mutant was detected (Fig. 2c, Fig. S3b). To search the sites of UEV-Vps23 ubiquitination, we performed a tandem mass spectrometry analysis to detect the ubiquitinated products, and confirmed that monoubiquitin moiety was covalently attached via an isopeptide bond at lysine residue 201 of the UEV-Vps23 (Fig. 2d). All these data exhibit that the UEV-Vps23 can undergo mono-ubiquitination through a eukaryotic-like ubiquitination cascade in Asgard archaea.

**Fig. 2.**
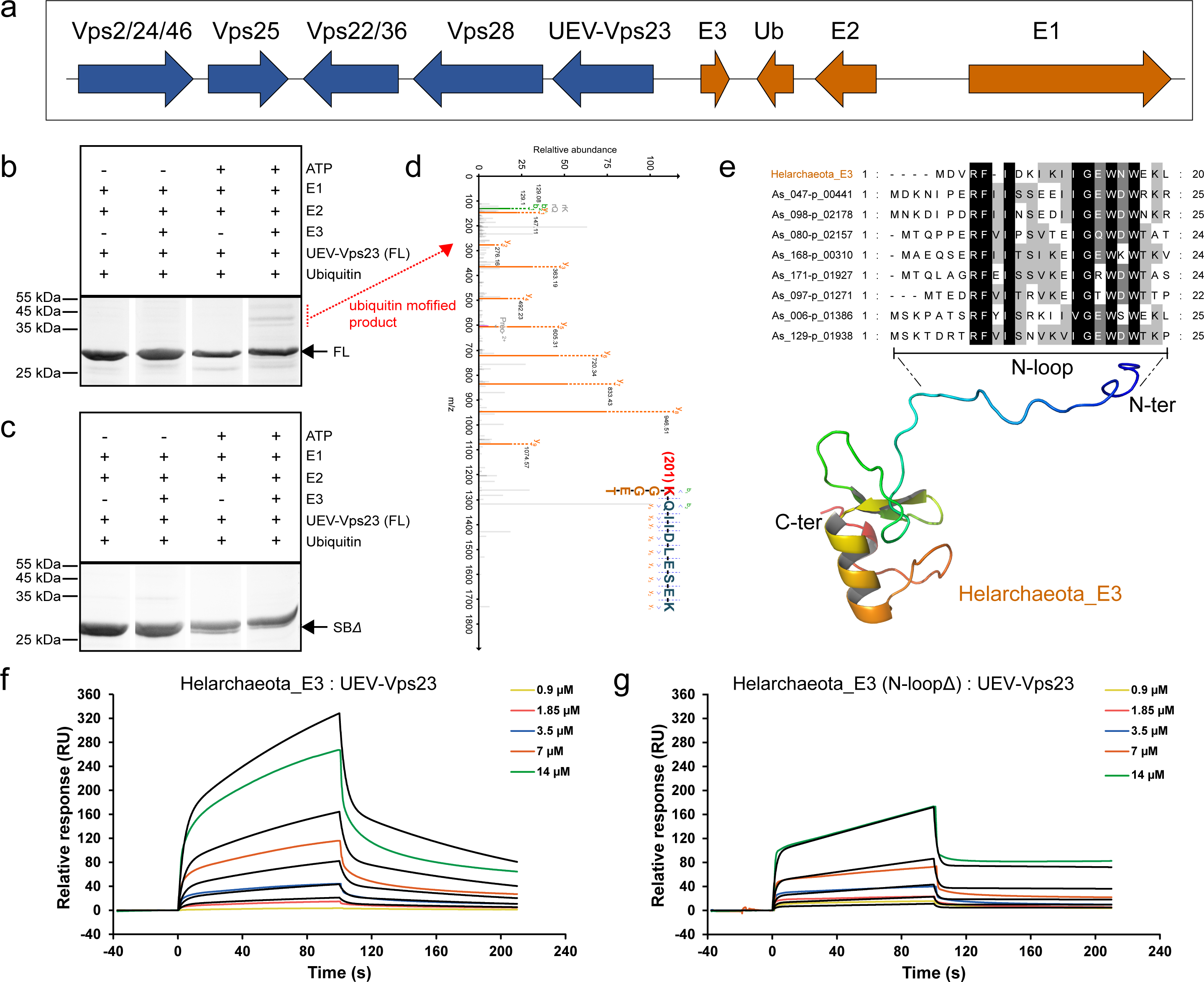
Ubiquitin modification of the UEV-Vps23 through a ubiquitination cascade in Helarchaeota. (a) Gene cluster of the ESCRT (colored blue) and ubiquitin system (colored orange) components in Helarchaeota. (b) Examining ubiquitin modification of the full-length UEV-Vps23 substrate *in vitro* ubiquitination reaction at 50 °C. The potential ubiquitin modified substrates that was further assayed by mass spectrometry are labeled by red dotted line. (c) Examining ubiquitin modification of the UEV-Vps23 (SB*Δ*) substrate *in vitro* ubiquitination reaction at 50 °C. (d) Tandem mass spectrometry analysis of ubiquitin modified full-length UEV-Vps23 *in vitro* ubiquitination reaction at 50 °C. Based on the mass spectra, the isopeptide bond between the C-terminal di-glycine motif (colored orange) and lysine residue 201 (colored red) at the steadiness box (colored blue) of the UEV-Vps23. (e) Identification of a conserved N-terminal loop region (N-loop) of the RING E3 using amino acid sequence alignment and structural modeling. The complete gels are shown in Fig. S3. (f) SPR analysis of interaction between UEV domain of UEV-Vps23 and full-length RING E3 (fitting a 2: 1 binding mode). (g) SPR analysis of interaction between UEV domain of UEV-Vps23 and full-length RING E3 (N-loop*Δ*) (fitting a 2: 1 binding mode).

The specific interaction between RING E3 and its target protein is critical for ubiquitination. In order to characterize the recognition model between the UEV-Vps23 and the RING E3, we aligned the amino acid sequences of this Asgard RING E3 and identified an evolutionarily conserved N-terminal region (Fig. 2e). Further structural modelling using ColabFold server (*43*) suggests that this region forms a loop structure (N-loop). To further determine if the N-loop is responsible for UEV recognition, we examined the interactions of Helarchaeia UEV with the full-length E3, and a truncated E3 that lacks N-loop (residues 1–19) using SPR assay. The results show that whereas UEV (residues 1–132) bound to both full-length E3 (*K*_D_ 2.57e^-4^ M; Fig. 2f, Table S2) and the truncated protein (*K*_D_ 8.14e^-3^ M, Fig. 2f), fitting a two-state reaction model, deletion of the E3 N-loop resulted in a 3.17-fold impaired binding capability. Collectively, these data confirm that the N-terminal loop region of the RING E3 facilitates UEV-Vps23 binding.

### Switch of UEV-Vps23 states in ESCRT and ubiquitin system interplay

It has been proposed that eukaryotic Vps23 may have an autoinhibited state, given that its UEV is able to bind the PTAP motif located between the stalk region and SB (*44*). Because of identification of a conserved PXXP motif in a similar position of UEV-Vps23, we sought to verify autoinhibition using SPR analysis (Fig. 3a). We immobilized the Helarchaeia UEV (residues 1– 132) on CM5 chip, and employed the Vps23 domain (residues 133–258) as an analyte. The SPR sensorgram indicates a clear response signal after injection of the Vps23 domain (10 μM), revealing a direct binding between the two domains (Fig. 3b). Followed by the Vps23 domain ligand, we injected Helarchaeia ubiquitin (10 μM) into the reaction, and observed an increase in RU response without the emergence of dissociation curve of bound Vps23 domain, indicating that the UEV binding regions of the Vps23 domain and ubiquitin are distinct. We therefore conclude that the autoinhibited UEV-Vps23 can recognize and capture ubiquitinated cargoes (Fig. 3c).

**Fig. 3.**
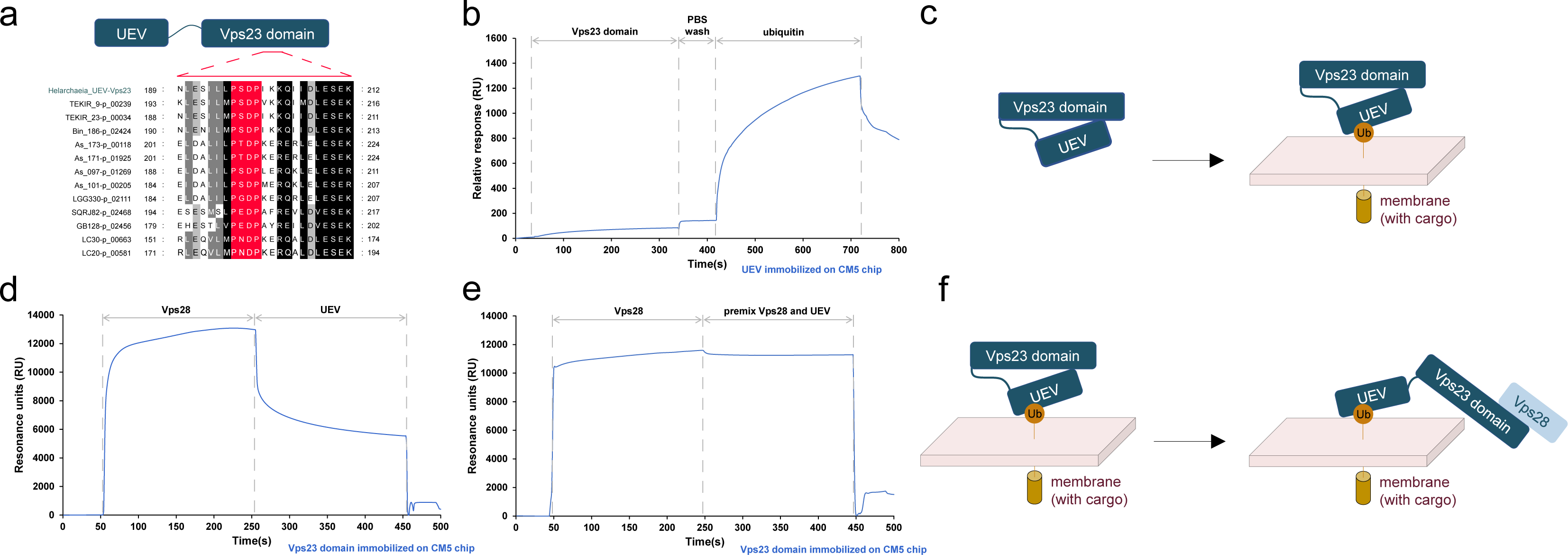
Switch of the UEV-Vps23 conformations in initiation of the ESCRT and ubiquitin system interplay. (a) The predicted PXXP motif for UEV-Vps23 autoinhibition are highlighted in red. Additional information on Asgard UEV-Vps23 can be found in Table S4 (b) SPR sensorgram obtained from the first injection of Vps23 domain and the second injection of ubiquitin over the UEV immobilized on CM5 chip. (c) Schematic representation of the ubiquitinated cargo targeting of the autoinhibited UEV-Vps23. (d) SPR sensorgram obtained from the first injection of Vps28 and the second injection of UEV over the Vps23 domain immobilized on CM5 chip. (e) SPR sensorgram obtained from the first injection of Vps28 and the second injection of premix of Vps28 and UEV over the Vps23 domain immobilized on CM5 chip. (f) Schematic representation of the Vps28 recruitment of the active UEV-Vps23. All the SPR tested UEV and Vps23 domain of UEV-Vps23, ubiquitin, and Vps28 are from the Helarchaeota genome (GCA_005191425_1).

To understand behaviors of the UEV-Vps23 in regulation of ESCRT and ubiquitin system interplay, we conducted an SPR binding exclusion experiment by injecting the Vps28 over the Vps23 domain that immobilized on CM5 chip, followed by injection of UEV domain. The SPR sensorgram displays that the adding the Vps28 (80 μM) led to a sharp increase in RU response, reflecting an interaction between the Vps28 and Vps23 domain, whereas the RU signal decreased after injection of UEV domain (80 μM), and this observation implies a blocking effect of the Vps28 on the binding of the UEV to Vps23 domain (Fig. 3d). To exclude the possibility that this decreasing RU signal was a result of dissociation between the Vps28 and Vps23 domain, we then carried out a corresponding SPR assay with the Vps23 domain on CM5 chip, using the Vps28 (40 μM) as the first injection, and a premix of UEV and Vps28 (40 μM: 40 μM) the second injection. The resulting sensorgram shows that the adding Vps28 induced a quick response equilibrium, whereas no further increasing RU signal was observed after injection of the premix, and hence demonstrates that the UEV was unable to bind to the Vps28-Vps23 domain complex (Fig. 3e). These data together reveal that the Vps28 and UEV compete for binding with Vps23 domain, and thus the autoinhibited UEV-Vps23 should be opened before recruitment of the downstream Vps28 (Fig. 3f).

Overall, the above SPR results display that the UEV of UEV-Vps23 can fold over the Vps23 domain, forming an autoinhibited state in Asgard cellular context. Molecular mechanisms, perhaps the assembly of Vps28, which transfer the UEV-Vps23 into open confirmation are critical for activating the putative ubiquitin-dependent sorting process.

### Evolutionary linkages among UEV-Vps23, eukaryotic Vps23, and archaeal CdvA

To explore the origin and evolution of the domains of UEV-Vps23, we collected E2/UEV and Vps23 domain-containing sequences in InterPro databases and related Asgard archaeal Clusters of Orthologous Genes in our previous work (*34*). As shown in the maximum-likelihood phylogenetic tree of E2/UEV domain (Fig. 4a), *Ca.* Thermoplasmatota E2 seems to have evolved earlier than Asgard and TACK archaeal E2 since the clade is placed at the deep branch of archaeal clade of UEV/E2. TACK archaeal E2 clade is nested with various Asgard archaeal groups, suggesting that TACK archaeal E2 were possibly horizontal transferred from Asgard archaea. Eukaryotic UEV domain sequences were diverged from a clade composed of mostly Heimdallarchaeia (cog.005829), and shared common ancestral relationship with Lokiarchaeia (cog.001811 and cog.000544). Eukaryotic UEV domain and eukaryotic E2 domain are likely evolved independently within the diverse Asgard archaeal E2/UEV clades.

**Fig. 4.**
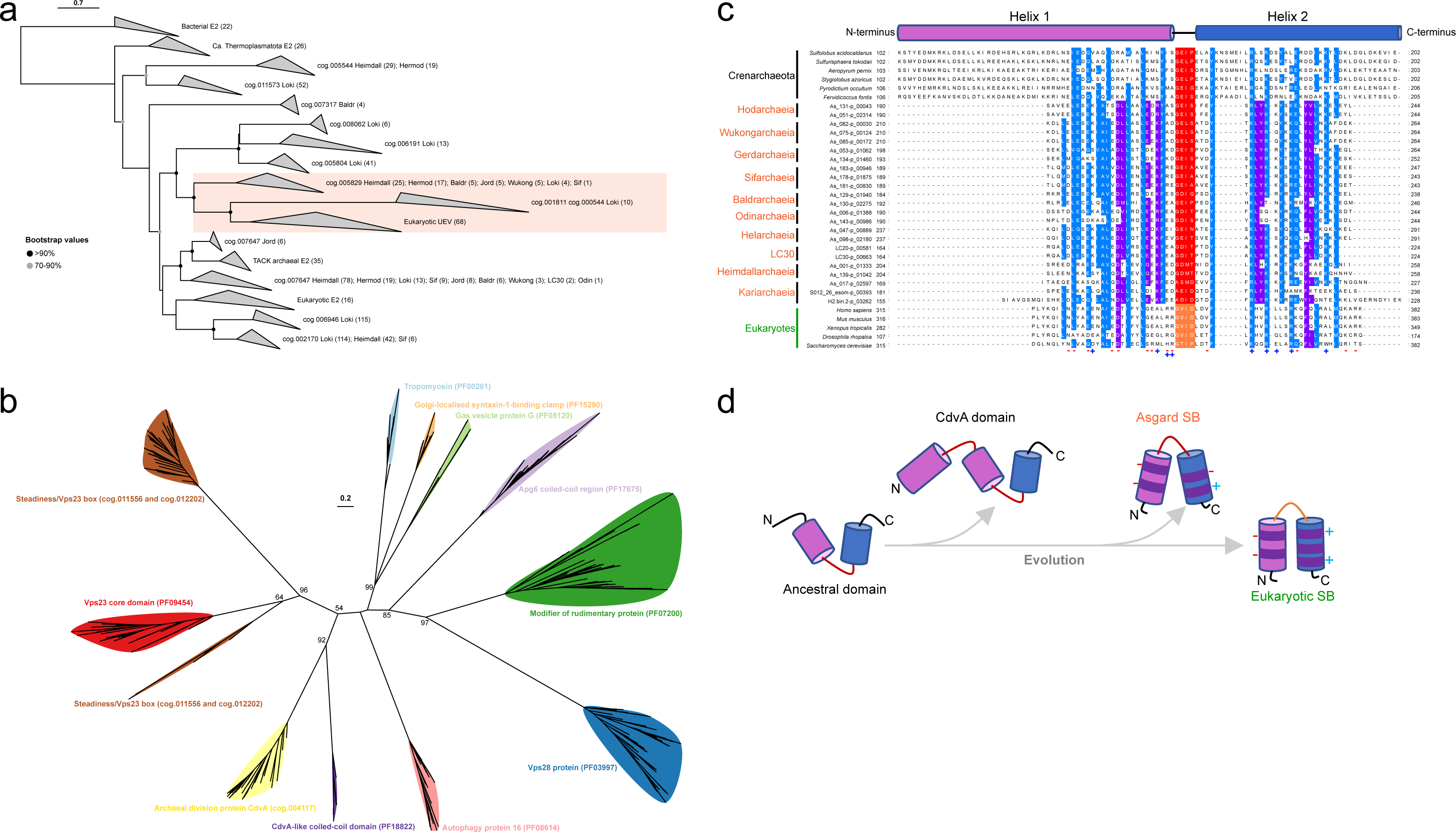
Evolutionary history of the Asgard UEV-Vps23. (a) The maximum-likelihood phylogenetic tree of E2/UEV domain was built from an alignment of 886 sequences with 277 amino acid residues. The red semi-transparent square highlighted the possible scenario that eukaryotic UEV originated from Asgard archaea. Bacterial A E2 homolog that is used as an outgroup as previously described (*14*). The numbers in brackets indicated the sequence quantities in the collapsed clades with taxonomy specificity. (b) the maximum-likelihood phylogenetic tree of steadiness box/Vps23 domain was built from an alignment of 421 sequences with 234 amino acid residues. Ultra-fast bootstrap values were shown. (c) Alignment of amino acid sequences of SB in UEV-Vps23 and Vps23, and of CdvA domains. The relatively conserved sequences of SB of the UEV-Vps23 and Vps23, and CdvA domain are shown explicitly. The distribution of positively (+) and negatively (-) charged residues in SB of the UEV-Vps23 and Vps23 are marked. Additional information on eukaryotic Vps23, and crenarchaeal CdvA domain, and Asgard UEV-Vps23 can be found in Table S3 and Table S4. (d) A scenario for the evolutionary history of SB in UEV-Vps23 and Vps23, and CdvA domain. The SB of UEV-Vps23 and Vps23, and CdvA domains were likely derived from an ancestral hairpin domain that was present in the common ancestor of TACK and Asgard archaea. The SB evolved as a result of increased charged residues, whereas the CdvA domain likely arose from the ancestral domain encoding gene duplication.

Next, we performed a phylogenetic analysis of steadiness box (SB)/Vps23 domain of the UEV-Vps23 and Vps23 as well as short helical segments of the Vps28, CdvA, Syntaphilin, Tropomyosin, ATG16, APG6, GvpG, and Modr. According to the unrooted maximum-likelihood phylogenetic tree (Fig. 4b), the SB/Vps23 domains seem to arise from the common ancestor of archaeal division protein CdvA and CdvA-like coiled-coil domains. Other short helical segment containing proteins evolved separately from SB/Vps23/CdvA domains.

In addition to the phylogenetic analyses, we found that UEV-Vps23 SB presents key residue and structural similarities with both the eukaryotic SB and the archaeal CdvA domains (Fig. 4c). In particular, the Asgard SB and the CdvA domain share the presence of a predicted loop region. To characterize those similarities, we further constructed the structure models of Asgard UEV-Vps23 and a CdvA from *Sulfolobus acidocaldarius*, using ColabFold server (*43*). The UEV-Vps23 SB adopts a helical hairpin structure that is structurally analogous to the SB of *S. cerevisiae* Vps23 (PDB: 2F6M), with root-mean-square deviation (RMSD) values in the range of 0.232–0.797 Å (Fig. S4). Importantly, we found that the SB structures of the UEV-Vps23 in Odin-(RMSD 0.379 Å), Gerd-(0.518 Å), LC30-(0.594 Å), and Borrarchaeia (1.015 Å) resemble part of the CdvA domain structure, despite that the two helices in the SB structures are shorter. Especially, the first helixes in both hairpins of Asgard and eukaryotic SB are enriched negative charged glutamate and aspartate residues, whereas the others are positively charged residues, including lysine and arginine. These charged residues probably provide stability to SB hairpin structure. Notably, we indeed observed the conserved loop region sequences in the CdvA domain, which might be attributed to gene duplication (Fig. S5). Together, these findings suggest that the Asgard Vps23 domains of UEV-Vps23 and eukaryotic Vps23, and CdvA domain likely descended from a common ancestral domain (Fig. 4d).

The results of phylogenetic analyses, sequence comparison, and structural modeling provide evidence for an ancestral relationship among UEV-Vps23, eukaryotic Vps23, and archaeal CdvA. It seems likely that the UEV evolved and fused with a CdvA-like Vps23 domain, alongside the emergence of ubiquitin system in Asgard archaea.

## Discussion

In this study, we describe an important role of a UEV-Vps23 family protein, that harbors UEV and Vps23 domains, in an interplay between ESCRT and ubiquitin system in Asgard archaea (Fig. S6). The strong interaction between the UEV and ubiquitin suggests that UEV-Vps23 senses and captures ubiquitinated cargoes, functionally coupling the ESCRT with ubiquitin system. The Vps23 domain was subjected to modification in a ubiquitination cascade, depending on its SB, implying that the UEV-Vps23 is post-transcriptionally modulated by the ubiquitin system. Furthermore, modulation of the interplay between ESCRT and ubiquitin system seems to be achieved through conformation switch between the autoinhibited and active forms of UEV-Vps23.

The SPR analysis indicates that Asgard UEV exhibits high affinity toward ubiquitin, compared to that of between eukaryotic UEV and ubiquitin. This distinction might be underpinned by the structural features of Asgard UEV, such as its apparent absence of β-tongue structure, an extended β-hairpin which is implicated in binding surface between eukaryotic UEV with ubiquitin (*36, 44*). Our functional reconstruction of a eukaryotic-like ubiquitination cascade in Asgard archaea is similar with the observation that a ubiquitin system of TACK archaeon *Caldiarchaeum subterraneum* ubiquitinated an artificial Rad50 substrate *in vitro* (*42*). It therefore speculates that the ubiquitin system had evolved in archaeal ancestor before the TACK and Asgard-eukaryotes split. It has been suggested that the SB ubiquitination holds an important role in control Vps23 expression in mammalian cells (*45*). It thus raises the possibility that the mono-ubiquitination of UEV-Vps23 is required for a proteasomal degradation pathway in Asgard cellular context. Combined with the recent finding that proteasome mediated ESCRT-III homologues degradation in *S. acidocaldarius*, our work suggests an evolutionarily conserved role for proteasome in control of ESCRT in TACK and Asgard archaea (*46*).

The results of SPR analysis offer an elaborate conformational switch for UEV-Vps23 in initiation of ESCRT and ubiquitin system interplay. It appears that Vps28 binding is essential and required for relieving UEV-Vps23 autoinhibited conformation. Since the Vps28-binding and ubiquitination sites are overlapped in the SB of UEV-Vps23, binding Vps28 may block the ubiquitination of UEV-Vps23 and thereby ensures stabilization of its active form. Likewise, in eukaryotic system, the Vps23 perhaps functions as a similar regulatory switch with dual modes of engagement in accurate control of ESCRT and ubiquitin system interplay, a molecular mechanism likely inherited from an archaeal ancestor.

The intriguing finding of our work is that the UEV-Vps23 exhibits sequence and structural traits common to both the eukaryotic Vps23 and crenarchaeal CdvA. The UEV of most UEV-Vps23 are fused with specific Vps23 domains that are likely related to archaeal CdvA domains. The divergence between such CdvA-like Vps23 domain and the eukaryotic-signature one seemingly occurred in Asgard archaea (*34*). These findings are compatible with common ancestral origin of CdvA and Vps23 homologs, which suggests, in a broader evolutionary context, that the ESCRT-I subunits emerged predating the radiation of the TACK and Asgard archaea.

An outstanding question is that what are the biological functions of the ESCRT and ubiquitin system interplay in Asgard archaea. It seems likely that the interplay points to the existence of a putative ubiquitin-sorting pathway in Asgard cells, which can be utilized for cargo sorting processes in internalization of cell surface receptors, and intracellular trafficking. Albeit the current observation of absence of eukaryotic-like endomembrane in such as Lokiarchaeota isolates, the ubiquitin sorting capability may be required for a directed long intracellular cargo transport in specific structures, such as long protrusions (*47, 48*).

In conclusion, we biochemically characterized the functionating of UEV-Vps23 in an ancestral form of ESCRT and ubiquitin system interplay which likely already emerged in the last Asgard common ancestor. Our finding of the evolutionary trajectory of Vps23 homologs shows that how origin and evolution of subdomains give rise to new molecular properties of entire protein, which consequently results in functional leap in the ESCRT evolution at cellular level. Overall, the presence of such ESCRT and ubiquitin system interplay raises the possibility that the Asgard archaea have a ubiquitin-sorting pathway, which might function as a scaffolding for a complex membrane trafficking system. Further molecular and cell biological research should be focused on this unprecedented membrane trafficking system to shed light on emergence of endomembrane system in eukaryogenesis.

## Materials and Methods

### Construction of E2, UEV, CdvA, and Vps23 domain-specific multiple sequence alignments

A total of 3447 eukaryotic Vps23 protein sequences were retrieved from InterPro database with a search for proteins with UEV domain (PF05743) and Vps23 core domain (PF09454) architecture; 64 eukaryotic ubiquitin-conjugating enzyme E2 protein sequences were obtained from PF00179 seed alignment; 86 bacterial E2 family protein sequences were retrieved with a search for proteins with Prokaryotic E2 family A (PF14457); 240 non-Asgard archaeal ubiquitin-conjugating enzyme domain containing protein sequences were retrieved with a search for proteins with UBC core (PF00179), on May 31^st^ 2023. Asgard archaeal sequences were obtained by search on Ubiquitin-conjugating enzyme E2 related AsCOGs (cog.011573, cog.007647, cog.006946, cog.002170, cog.005544, cog.007313, cog.000544, cog.001811, cog.005829, cog008062, cog.005804 and cog.006191) with Hidden Markov Model by a 1E-10 e-value cutoff. E2/UEV domain region coordinates of these sequences were extracted according to 1) their annotation information provided by UniProt database; 2) results from hhsearch on Pfam v35 database. For each cluster of putative orthologous genes, outlier sequences potentially derived from misannotation or inadequate cutoff were identified by CIAlign package (*49, 50*) with a “--remove_divergent_minperc 0.5” flag. The clean set of multiple sequence alignment of E2/UEV domain sequences was constructed using a high accuracy MAFFT mode L-INS-I (*51*). The alignment was then trimmed using CIAlign (settings:--remove_insertions--remove_short-- remove_min_length 50) to remove insertions which are not present in the majority of sequences. The alignment was used to reconstruct a phylogenetic tree using IQ-TREE (*52*) with best-fit evolutionary model and site rates and 1000 times ultrafast bootstrap approximation (-m LG+F+R7-B 1000)(*53*).

The evolutionary analysis of steadiness box domain was conducted in a similar way. Steadiness box region containing protein sequences were retrieved from InterPro database with a search for proteins with CdvA-like coiled-coil domain (PF18822) and Vps23 core domain (PF09454) architecture. Asgard archaeal sequences were obtained by search on archaeal division protein CdvA (cog.004117) and Steadiness/Vps23 box (cog.012202 and cog.011556) with Hidden Markov Model by a 1E-10 e-value cutoff. Other helical protein families containing short helical segments and a loop were added to the phylogenetic analysis, i.e., Syntaphilin (PF15290); Tropomyosin (PF00261); ATG16 (PF08614); APG6 (PF17675); GvpG (PF05120); Modr (PF07200) and Vps28 (PF03997). The best-fit evolutionary model for steadiness box containing protein was “LG+F+G4”. The visualization of the phylogenetic tree was realized with tvBOT (*54*). Taxonomy information of archaeal genomes was obtained from GTDB RS214 (*55*).

### Structural modelling

The three-dimensional structures of the full length of UEV-Vps23, Helarchaeota RING E3, and *Sulfolobus acidocaldarius* CdvA (UniProtKB accession number Q4J923) were constructed using ColabFold server (*43*). The multiple sequence alignment model, model type, pair mode, and the number of recycle was set as “UniRef+Environmental”, “auto”, “unpaired+paired”, and “3”, respectively. Each protein sequence was generated five model structures, the one of which with the highest IDDT score was used in this paper. The structural analyses were carried out by using PyMOL software (http://www.pymol.org).

### Protein expression and purification

The coding sequences for proteins and domains used here were codon optimized and synthesized by Tsingke Biotech Co., Ltd (China), and, respectively, cloned into a pCold-II vector (Takara Bio Co Ltd., Japan) containing an N-terminal His tag. These recombinant plasmids were transformed into *Escherichia coli* BL21 (Takara Bio Co Ltd, Japan) and the transformed cells were cultured at 37°C until the OD_600_ reached at 0.6–0.8, before inducing by 0.2–0.5 mM isopropyl-d-1-thiogalactopyranodside at 15 °C for 18 h. The cells pellets were harvested by centrifugation, and resuspended in 20 ml Binding Buffer (20 mM phosphate buffer solution (pH 7.4), 300 mM NaCl, 50 mM imidazole, 1 mM dithiothreitol, and 1×Protease Inhibitor Cocktail (BBI Co., Ltd, China)) before ultrasonic decomposition. Then, the proteins and domains were purified by ÄKTA Go System (GE Healthcare, USA) with Ni-NTA affinity chromatography column (HisTrap HP, GE Healthcare, USA) and size exclusion chromatography column (Superdex 200 Increase 10/300 GL, GE Healthcare, USA). Finally, the purified proteins and domains were pooled and concentrated to 0.2–1 ml in PBS buffer by Amicon Ultra-15 (Millipore, USA). A BCA Protein Assay Kit (Beytotime Bio Co Ltd., China) was applied to assay the concentrations of the proteins and domains. The *Homo sapiens* ubiquitin was purchased from Sigma-Aldrich Chemical corporation (USA) and was suspended in 100 mM phosphate buffer solution (pH 7.4).

### Ubiquitination reaction

For the E1 and E2 ubiquitination assay, 50 μg E1 protein, 50 μg E2 protein and 50 μg mature ubiquitin in the presence or absence 3.3 mM ATP in 200 μl reaction buffer (20 mM Tris-HCl pH 8.0, 150 mM NaCl, 5% glycerol, 5 mM MgCl_2_, 1 mM dithiothreitol) for 30 min at 50 °C and 60 °C, respectively. For the E1, E2, and RING E3 cascade assay, 50 μg mature ubiquitin, 10 μg E1, 10 μg E2, 50 μg RING E3, and 30 μg Helarchaeota UEV-Vps23 (or UEV-Vps23 SB*Δ*) substrate incubated in 200 μl reaction buffer for 60 min at 60 °C in the presence or absence of 2.5 mM ATP (with addition of fresh ATP each 15 min). The resulting products were analyzed by SDS-PAGE, followed by Coomassie staining.

### Surface plasmon resonance assay

Surface plasmon resonance assay was performed using Biacore 8k (GE Healthcare, USA). The purified proteins or domains were immobilized on CM5 chip (GE Healthcare, USA) at response unit 2914.6, 2719.4, 11358.5, 2587.2 and 2797.0, respectively, using the standard primary amine coupling method as described previously (*56*). The purified ligands in 100 mM phosphate buffer solution (pH 7.4) were run at a flow rate of 30 µl/min at 25 °C. The resulting data were calculated using BIA evaluation software (GE Healthcare, USA).

### Mass spectrometry analysis

The ubiquitinated products were subjected to digestion by trypsin at 37 °C for 14–16 h, and were then desalted using Waters solid phase extraction cartridges before LC-MS/MS analysis. The LC-MS/MS experiments were performed using Thermo Scientific UltiMate™ 3000 Binary Rapid Separation System and Thermo Scientific EASY-nLC™ 1200 System. The separation of peptides was achieved through reversed-phase high-pressure liquid chromatography, employing an Agilent ZORBAX 300 Extend-C18 column (3.5 μm, 4.6 ×150 mm) at a flow rate of 0.3 ml/min. The separated peptides were collected and injected into a self-loading C18 column (100 μm i.d., 1.8 μm particle size) operating at a flow rate of 300 nl/min.

The peptides were separated using the following gradient: from 0 to 103 min, solvent B (98% ACN, 0.1% FA) was linearly increased from 4% to 27%; from 103 to 111 min, solvent B was increased from 27% to 40%; from 111 to 113 min, solvent B was increased from 40% to 90%; from 113 to 120 min, the system operated with 90% solvent B. The separated peptides were ionized by a nano-ElectroSpray Ionization (ESI) and transferred to an Orbitrap Exploris™ 480 mass spectrometer (Thermo Fisher Scientific, San Jose, CA) for detection in DDA (Data dependent Acquisition) mode. The parameter for mass spectrometer were set as follows: ion source voltage was 2.2 KV; primary mass spectrometry scan range was 350–1,500 m/z with a resolution of 60,000. The normalized AGC Target was 300%, and maximum ion injection time was 20 ms; The secondary mass spectrometry fragmentation mode was HCD. The fragmentation energy was set at 32% and a resolution of 15,000. The dynamic exclusion time was 60 s. The starting m/z of secondary mass spectrometry was fixed to 110 and parent ion screening condition for secondary fragmentation was charge 2+ to 6+. The Normalized AGC Target was set at standard, and the maximum ion injection time was 22 ms.

MaxQuant software (version 2.1.4.0) was used for the analysis of label-free MS/MS data. The following settings were utilized: type: standard; enzyme: Trypsin/P; maximum missed cleavages: 2; fixed modification: carbamidomethyl (C); variable modifications: oxidation (M) and acetyl (protein N-term); precursor mass tolerance: 20 ppm; fragment mass tolerance: 0.05 Da; The MS/MS data were searched against a custom database containing UEV-Vps23 and ubiquitin-GG (di-glycine motif). To ensure reliable results, a false discovery rate threshold of 1% was applied to both peptide spectrum match and protein levels. Protein from contaminant or reverse was removed.

## Acknowledgements

We appreciate the constructive suggestions on the work from Kira S. Makarova and Eugene V. Koonin (National Center for Biotechnology Information, National Library of Medicine, Bethesda, USA), and the technical support for surface plasmon resonance assay by Mengsi Sun (Shenzhen Bay Laboratory, Shenzhen, China).

## Author contributions

Z.Y.L. and M.L. conceived and designed the experiments. S.Y.Z. and R.Y.X. performed the experiments. Z.Y.L. and Y.L. performed the bioinformatics analyses. Z.Y.L., S.Y.Z., and Y.L. analyzed the data. Z.Y.L., S.Y.Z., Y.L., and M.L. wrote the paper, and all authors edited and approved the paper.

## Data and materials availability

All data needed to evaluate the conclusions in the paper are present in the paper and/or the Supplementary Materials.

Additional data related to this paper may be requested from the authors.

## References

1. J. B. Dacks, A. A. Peden, M. C. Field, Evolution of specificity in the eukaryotic endomembrane system. Int J Biochem Cell Biol 41, 330–340 (2009).

2. W. Martin, E. V. Koonin, Introns and the origin of nucleus-cytosol compartmentalization. Nature 440, 41–45 (2006).

3. H. M. McBride, Mitochondria and endomembrane origins. Curr Biol 28, R367–R372 (2018).

4. J. B. Dacks, M. C. Field, Evolutionary origins and specialisation of membrane transport. Curr Opin Cell Biol 53, 70–76 (2018).

5. M. A. OMalley, M. M. Leg er, J. G. Wideman, I. Ruiz-Trillo, Concepts of the last eukaryotic common ancestor. Nat Ecol Evol 3, 338–344 (2019).

6. W. M. Henne, N. J. Buchkovich, S. D. Emr, The ESCRT pathway. Dev Cell 21, 77–91 (2011).

7. S. B. Shields, R. C. Piper, How ubiquitin functions with ESCRTs. Traffic 12, 1306–1317 (2011).

8. M. Vietri, M. Radulovic, H. Stenmark, The many functions of ESCRTs. Nat Rev Mol Cell Biol 21, 25–42 (2020).

9. H. Teo, D. B. Veprintsev, R. L. Williams, Structural insights into endosomal sorting complex required for transport (ESCRT-I) recognition of ubiquitinated proteins. Journal of Biological Chemistry 279, 28689–28696 (2004).

10. H. Teo et al., ESCRT-I core and ESCRT-II GLUE domain structures reveal role for GLUE in linking to ESCRT-I and membranes. Cell 125, 99–111 (2006).

11. D. J. Katzmann, C. J. Stefan, M. Babst, S. D. Emr, Vps27 recruits ESCRT machinery to endosomes during MVB sorting. J Cell Biol 162, 413–423 (2003).

12. T. Wollert, C. Wunder, J. Lippincott-Schwartz, J. H. Hurley, Membrane scission by the ESCRT-III complex. Nature 458, 172–177 (2009).

13. D. J. Katzmann, M. Babst, S. D. Emr, Ubiquitin-dependent sorting into the multivesicular body pathway requires the function of a conserved endosomal protein sorting complex, ESCRT-I. Cell 106, 145–155 (2001).

14. A. M. Burroughs, M. Jaffee, L. M. Iyer, L. Aravind, Anatomy of the E2 ligase fold: implications for enzymology and evolution of ubiquitin/Ub-like protein conjugation. J Struct Biol 162, 205–218 (2008).

15. O. Pornillos et al., Structure and functional interactions of the Tsg101 UEV domain. EMBO J 21, 2397–2406 (2002).

16. M. S. Kostelansky et al., Structural and functional organization of the ESCRT-I trafficking complex. Cell 125, 113–126 (2006).

17. J. T. White, D. Toptygin, R. Cohen, N. Murphy, V. J. Hilser, Structural stability of the coiled-coil domain of tumor susceptibility gene (TSG)-101. Biochemistry 56, 4646–4655 (2017).

18. B. McDonald, J. Martin-Serrano, Regulation of Tsg101 expression by the steadiness box: a role of Tsg101-associated ligase. Mol Biol Cell 19, 754–763 (2008).

19. Z. Erpapazoglou et al., A dual role for K63-linked ubiquitin chains in multivesicular body biogenesis and cargo sorting. Mol Biol Cell 23, 2170–2183 (2012).

20. B. Y. Kim, J. A. Olzmann, G. S. Barsh, L.-S. Chin, L. Li, Spongiform neurodegeneration-associated E3 ligase Mahogunin ubiquitylates TSG101 and regulates endosomal trafficking. Molecular biology of the cell 18, 1129–1142 (2007).

21. B. Korbei, Ubiquitination of the ubiquitin-binding machinery: how early ESCRT components are controlled. Essays in Biochemistry 66, 169–177 (2022).

22. K. S. Makarova, N. Yutin, S. D. Bell, E. V. Koonin, Evolution of diverse cell division and vesicle formation systems in Archaea. Nat Rev Microbiol 8, 731–741 (2010).

23. E. A. Pelve et al., Cdv-based cell division and cell cycle organization in the thaumarchaeon Nitrosopumilus maritimus. Molecular microbiology 82, 555–566 (2011).

24. J. Liu et al., Functional assignment of multiple ESCRT-III homologs in cell division and budding in Sulfolobus islandicus. Mol Microbiol 105, 540–553 (2017).

25. J. Liu et al., Archaeal extracellular vesicles are produced in an ESCRT-dependent manner and promote gene transfer and nutrient cycling in extreme environments. ISME J 15, 2892–2905 (2021).

26. L. Guy, T. J. Ettema, The archaeal TACK superphylum and the origin of euka ryotes. Trends Microbiol 19, 580–587 (2011).

27. R. Y. Samson et al., Molecular and structural basis of ESCRT-III recruitment to membranes during archaeal cell division. Mol Cell 41, 186–196 (2011).

28. Y. Caspi, C. Dekker, Dividing the Archaeal Way: The Ancient Cdv Cell-Division Machinery. Front Microbiol 9, 174 (2018).

29. A. Spang et al., Complex archaea that bridge the gap between prokaryotes and eukaryotes. Nature 521, 173–179 (2015).

30. K. Zaremba-Niedzwiedzka et al., Asgard archaea illuminate the origin of eukaryotic cellular complexity. Nature 541, 353–358 (2017).

31. K. W. Seitz et al., Asgard archaea capable of anaerobic hydrocarbon cycling. Nat Commun 10, 1822 (2019).

32. M. Cai et al., Diverse Asgard archaea including the novel phylum Gerdarchaeota participate in organic matter degradation. Science China Life Sciences 63, 886–897 (2020).

33. P. A. Bulzu et al., Casting light on Asgardarchaeota metabolism in a sunlit microoxic niche. Nat Microbiol 4, 1129–1137 (2019).

34. Y. Liu et al., Expanded diversity of Asgard archaea and their relationships with eukaryotes. Nature 593, 553–557 (2021).

35. Z. Lu et al., Coevolution of Eukaryote-like Vps4 and ESCRT-III Subunits in the Asgard Archaea. mBio 11, (2020).

36. T. Hatano et al., Asgard archaea shed light on the evolutionary origins of the eukaryotic ubiquitin-ESCRT machinery. Nat Commun 13, 3398 (2022).

37. M. Chakroun, N. Banyuls, Y. Bel, B. Escriche, J. Ferre, Bacterial Vegetative Insecticidal Proteins (Vip) from Entomopathogenic Bacteria. Microbiol Mol Biol Rev 80, 329–350 (2016).

38. H. J. Tran, M. D. Allen, J. Lowe, M. Bycroft, Structure of the Jab1/MPN domain and its implications for proteasome function. Biochemistry 42, 11460–11465 (2003).

39. S. Shaik, H. Pandey, S. K. Thirumalasetti, N. Nakamura, Characteristics and Functions of the Yip1 Domain Family (YIPF), Multi-Span Transmembrane Proteins Mainly Localized to the Golgi Apparatus. Front Cell Dev Biol 7, 130 (2019).

40. M. Heidtman, C. Z. Chen, R. N. Collins, C. Barlowe, Yos1p is a novel subunit of the Yip1p– Yif1p complex and is required for transport between the endoplasmic reticulum and the Golgi complex. Molecular biology of the cell 16, 1673–1683 (2005).

41. L. Hicke, H. L. Schubert, C. P. Hill, Ubiquitin-binding domains. Nature reviews Molecular cell biology 6, 610–621 (2005).

42. R. Hennell James et al., Functional reconstruction of a eukaryotic-like E1/E2/(RING) E3 ubiquitylation cascade from an uncultured archaeon. Nat Commun 8, 1120 (2017).

43. M. Mirdita et al., ColabFold: making protein folding accessible to all. Nature Methods, 1–4 (2022).

44. O. Pornillos et al., Structure and functional interactions of the Tsg101 UEV domain. The EMBO journal 21, 2397–2406 (2002).

45. B. McDonald, J. Martin-Serrano, Regulation of Tsg101 expression by the steadiness box: a role of Tsg101-associated ligase. Molecular biology of the cell 19, 754–763 (2008).

46. G. Tarrason Risa et al., The proteasome controls ESCRT-III-mediated cell division in an archaeon. Science 369, (2020).

47. H. Imachi et al., Isolation of an archaeon at the prokaryote–eukaryote interface. Nature 577, 519–525 (2020).

48. T. Rodrigues-Oliveira et al., Actin cytoskeleton and complex cell architecture in an Asgard archaeon. Nature, (2022).

49. C. Tumescheit, A. E. Firth, K. Brown, CIAlign: A highly customisable command line tool to clean, interpret and visualise multiple sequence alignments. PeerJ 10, e12983 (2022).

50. A. Comte et al., PhylteR: efficient identification of outlier sequences in phylogenomic datasets. bioRxiv, 2023.2002.2002.526888 (2023).

51. K. Katoh, D. M. Standley, MAFFT multiple sequence alignment software version 7: improvements in performance and usability. Mol Biol Evol 30, 772–780 (2013).

52. L. T. Nguyen, H. A. Schmidt, A. von Haeseler, B. Q. Minh, IQ-TREE: a fast and effective stochastic algorithm for estimating maximum-likelihood phylogenies. Mol Biol Evol 32, 268–274 (2015).

53. D. T. Hoang, O. Chernomor, A. von Haeseler, B. Q. Minh, L. S. Vinh, UFBoot2: Improving the Ultrafast Bootstrap Approximation. Mol Biol Evol 35, 518–522 (2018).

54. J. Xie et al., Tree Visualization By One Table (tvBOT): a web application for visualizing, modifying and annotating phylogenetic trees. Nucleic Acids Research, gkad359 (2023).

55. D. H. Parks et al., GTDB: an ongoing census of bacterial and archaeal diversity through a phylogenetically consistent, rank normalized and complete genome-based taxonomy. Nucleic acids research 50, D785–D794 (2022).

56. M. Ferrari et al., Characterization of a Novel ACE2-Based Therapeutic with Enhanced Rather than Reduced Activity against SARS-CoV-2 Variants. Journal of Virology 95, e00685–00621 (2021).

