## Supplementary material for "Origin of eukaryotic-like Vps23 shapes an ancient functional interplay between ESCRT and ubiquitin system in Asgard archaea": supplementary materials.pdf

For papers with three or more authors: Zhongyi Lu *et al.*

**This PDF file includes:**

Figs. S1 to S6
Tables S1 to S4

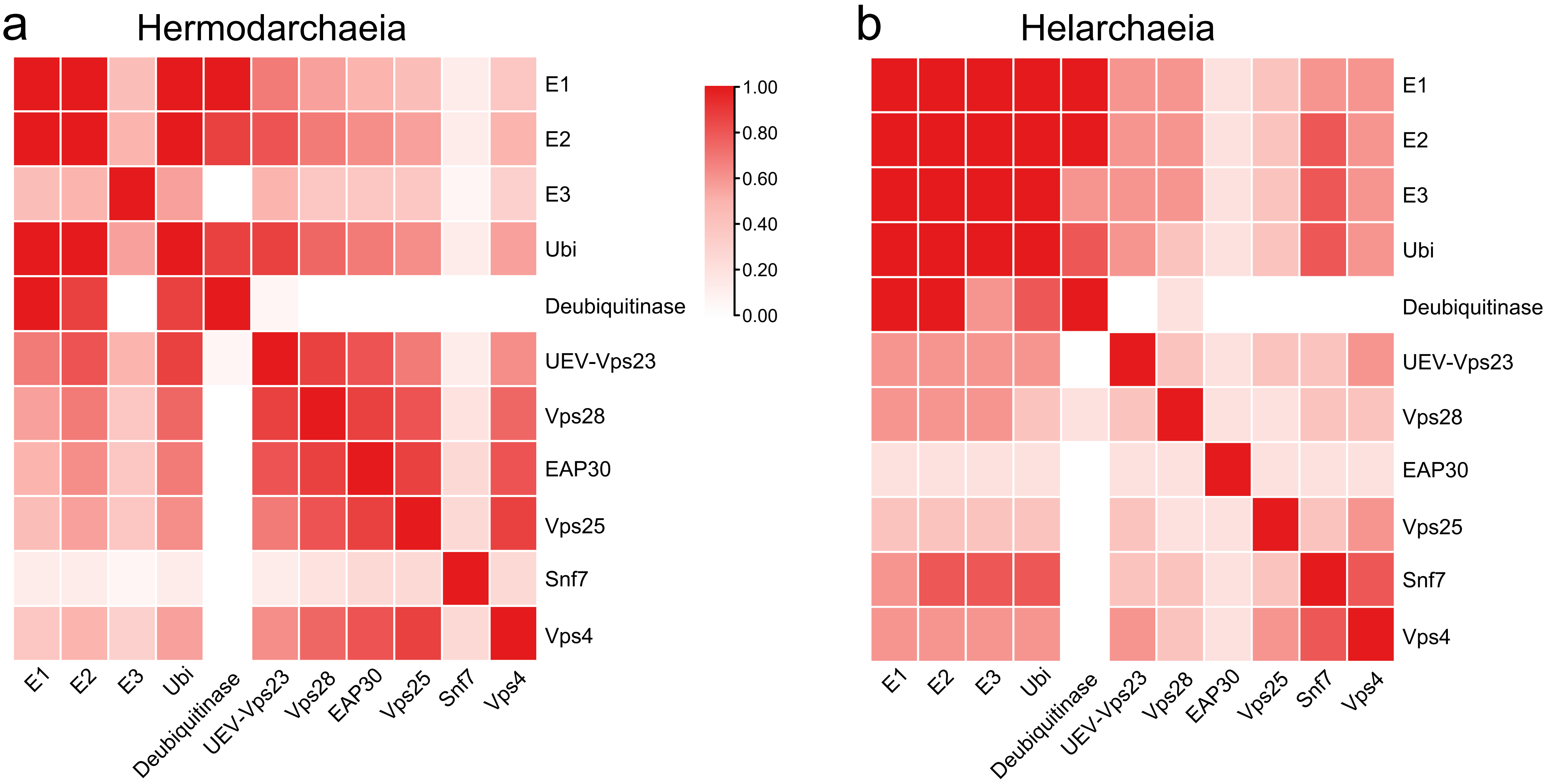


**Fig. S1.**

Colocation of genes encoding ESCRT and ubiquitin system components in Hermod- and Helarchaeia. A color gradient faction of genomes in which a pair of genes was found to cluster (a region of < 10kb).

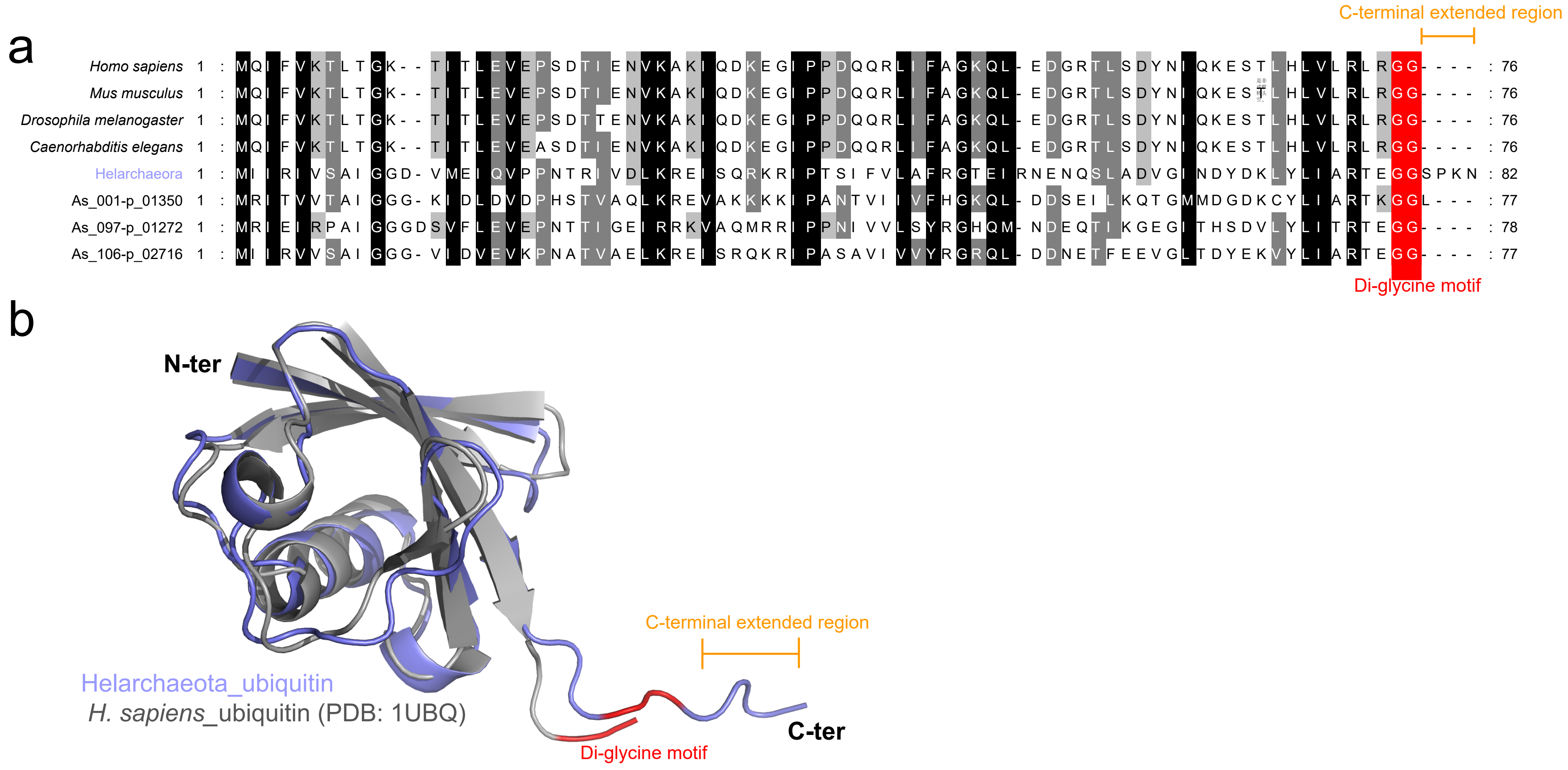


**Fig. S2.**

**Amino acid sequence analysis and structural modelling of the Asgard and eukaryotic ubiquitin.** (a) Alignment of amino acid sequences of the Asgard and eukaryotic ubiquitin. The ubiquitin sequences of *Homo sapiens* (PDB accession number 1UBQ), *Mus musculus* (PDB accession number 3VHT), *Drosophila melanogaster* (NCBI accession number AAA29000.1), and *Caenorhabditis elegans* (NCBI accession number NP 499695.1) are presented. Additional information on Asgard ubiquitin can be found in Table S4. The conserved di-glycine motif and C-terminal extended region of the ubiquitin are marked, respectively. (b) Structural superposition of the Asgard (purple) and *Homo sapiens* (grey) ubiquitin with a RSMD value (1.095 Å).

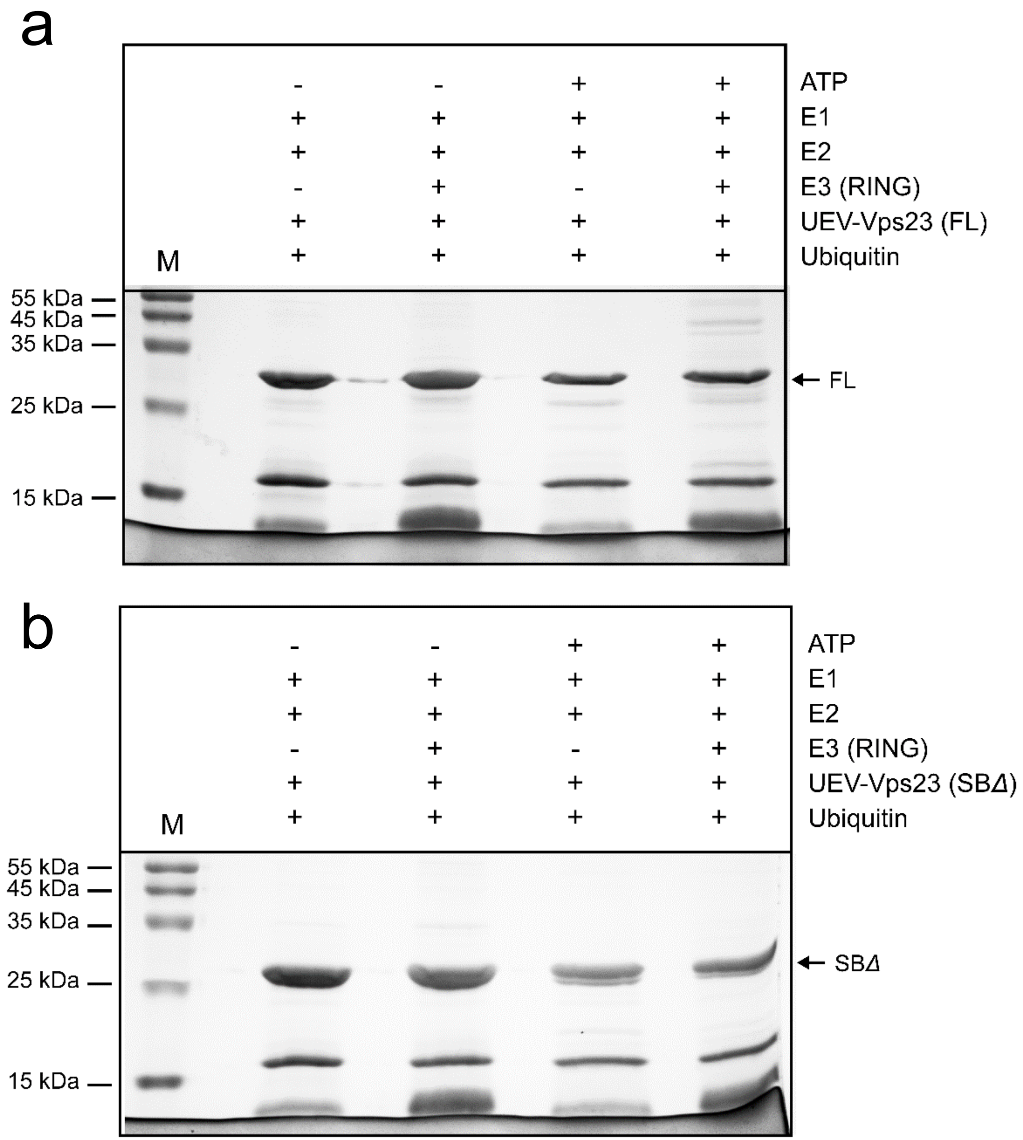


**Figure S3.**

**Full gels used for generation of main paper**. Shown are the complete gels used in Fig. 2b, upper (a) and down (b).

a

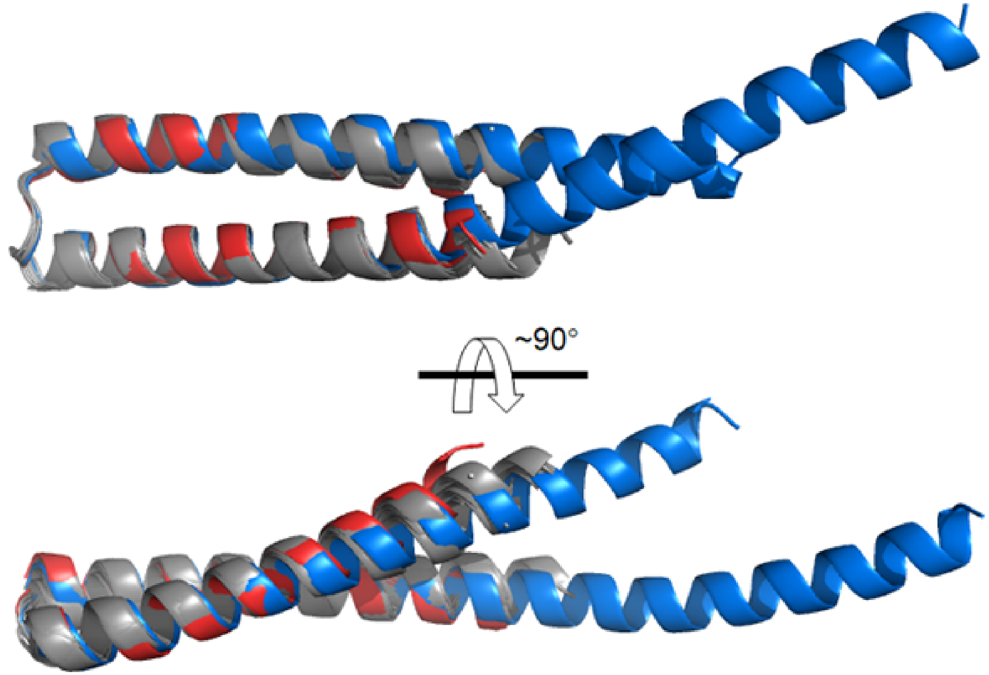


b

| | *S. cerevisiae* | *S. solfataricus* |
|---|---|---|
| Odinarchaeota | 0.64 | 0.379 |
| Wukongarchaeota | 0.554 | 4.634 |
| LC30 | 0.68 | 0.594 |
| Helarchaeota | 0.643 | 10.415 |
| Gerdarchaeota | 0.797 | 0.518 |
| Baldrarchaeota | 0.628 | 5.274 |
| Borrarchaeota | 0.677 | 1.015 |
| Hodarchaeota | 0.678 | 8.829 |
| Kariarchaeota | 0.232 | 4.944 |

**Fig. S4.**

**Structural characterization of the UEV-Vps23 SB.** Structural superposition (a) of the UEV-Vps23 SB (grey), the *Saccharomyces cerevisiae* Vps23 SB (red), and part of *Sulfolobus acidocaldarius* CdvA domain (blue) and the RSMD values (b). The RMSD values (Å) were calculated by using PyMOL software with 5 cycles of refinement for structural comparison of *Saccharomyces cerevisiae* Vps23 (PDB: 2F6M) and Asgard UEV-Vps23 structural models (Odinarchaeia, residues 190-251; Helarchaeia, 197-258; Gerdarchaeia, 194-252; Baldrarchaeia, 180-242; Wukongarchaeia, 206-271; LC30, 185-248; Borrarchaeia, 182-245; Hodarchaeia, 185-244; Kariarchaeia, 181-237), and part of *Sulfolobus acidocaldarius* CdvA domain (residues 131-194).

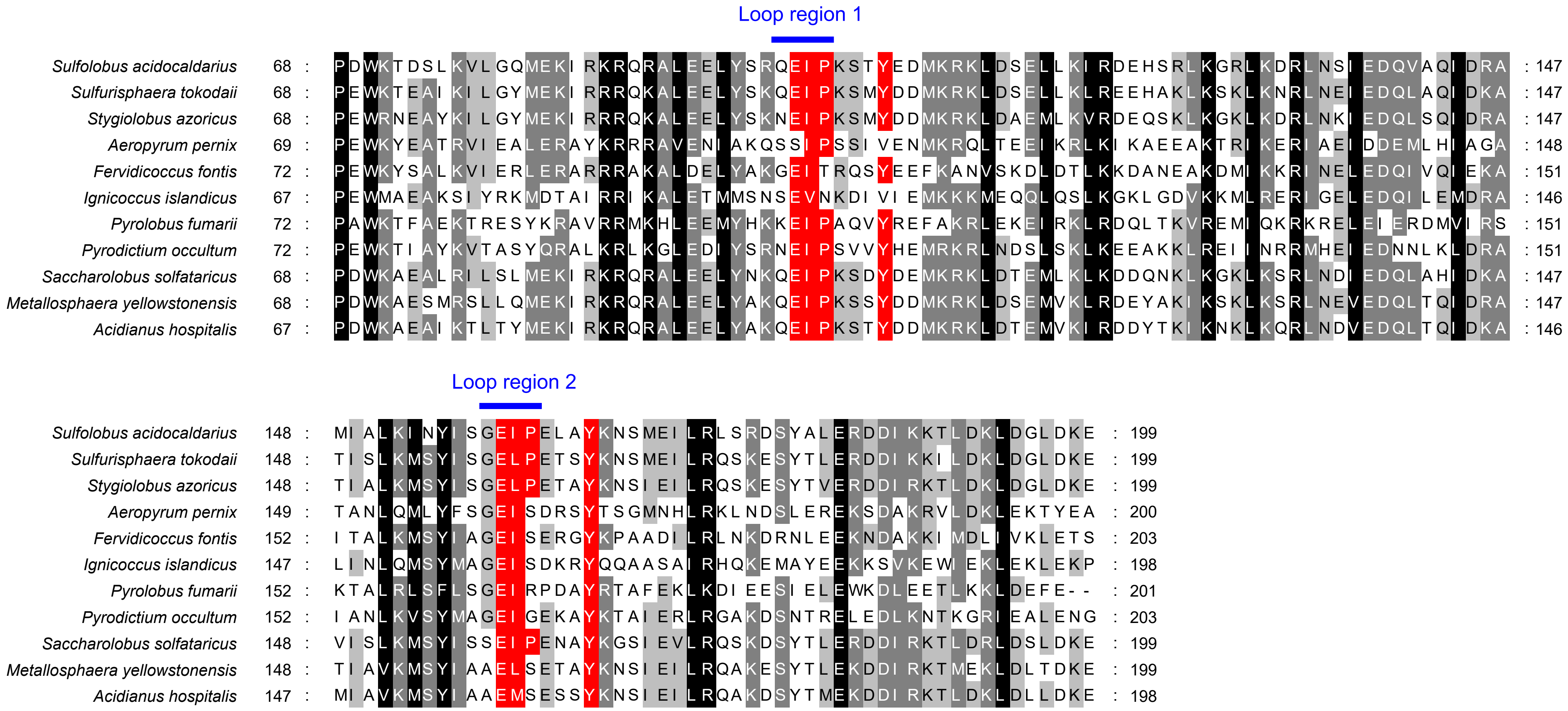


**Fig. S5.**

**Alignment amino acid sequences of CdvA domains.** The conserved amino acid residues around loop regions of CdvA domains are highlighted in red.

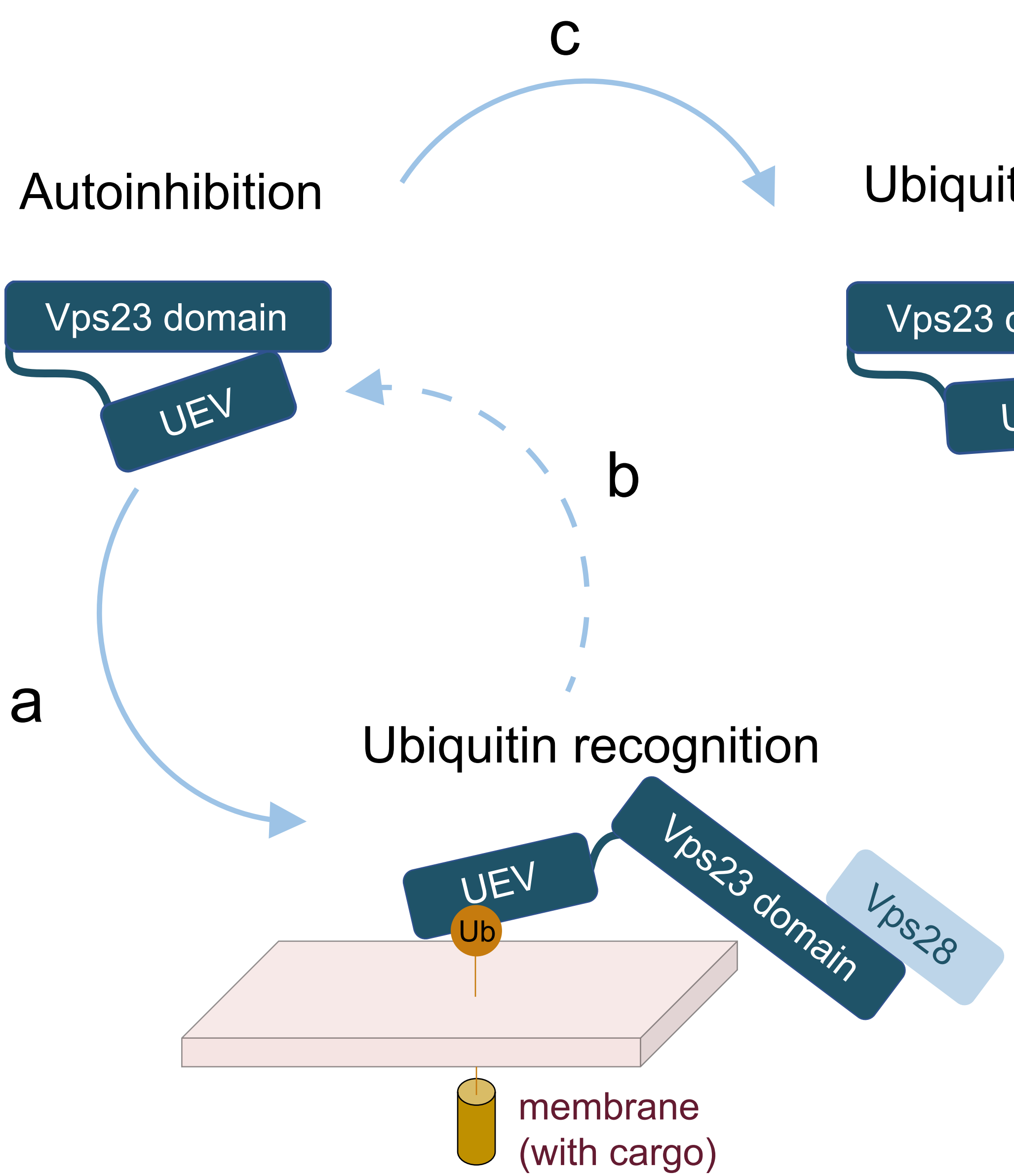


**Fig. S6.**

**Schematic model of key roles of the UEV-Vps23 in functionally connecting the ESCRT and ubiquitin system in Asgard archaea.** In Asgard cellular context, the free UEV-Vps23 has autoinhibition state before targeting to the ubiquitinated cargoes via the UEV (a). The autoinhibited UEV-Vps23 is activated and followed by recruitment of the downstream Vps28, conducting a ubiquitin-dependent sorting process (b). We proposed that the UEV-Vps23 releases from the ubiquitinated cargoes, ending up with the autoinhibited state. The SB of UEV-Vps23 can be ubiquitinated, through which the UEV-Vps23 is strictly controlled by the ubiquitin system (c).

### Table S1.

### Surface plasmon resonance analysis of interactions of Asgard UEV with Asgard ubiquitin and *Homo sapiens* ubiquitin.

| Immobilized on CM5 chip | Injection | Concentration (μM) | $K_D$ (M) |
|---|---|---|---|
| *S. cerevisiae* Vps23 $UEV_{1–171}$ | *H. sapiens* ubiquitin | 50, 25, 12.5, 6.25, 3.13, 1.56 | $3.79e^{-2}$ |
| Odin UEV-Vps23 $UEV_{1–130}$ | Odin ubiquitin | 50, 25, 12.5, 6.25, 3.13, 1.56 | $6.74e^{-7}$ |
| Odin UEV-Vps23 $UEV_{1–130}$ | *H. sapiens* ubiquitin | 50, 25, 12.5, 6.25, 3.13, 1.56 | $1.05e^{-4}$ |
| Wukong UEV-Vps23 $UEV_{1–138}$ | Wukong ubiquitin | 50, 25, 12.5, 6.25, 3.13, 1.56 | $3.34e^{-7}$ |
| Wukong UEV-Vps23 $UEV_{1–138}$ | *H. sapiens* ubiquitin | 50, 25, 12.5, 6.25, 3.13, 1.56 | $4.97e^{-5}$ |
| Heimdall UEV-Vps28 $UEV_{1–133}$ | Heimdall ubiquitin | 50, 25, 12.5, 6.25, 3.13, 1.56 | $4.39e^{-6}$ |
| Heimdall UEV-Vps28 $UEV_{1–133}$ | *H. sapiens* ubiquitin | 50, 25, 12.5, 6.25, 3.13, 1.56 | $4.53e^{-6}$ |
| Heimdall UEV-Vps23 $UEV_{1–131}$ | Heimdall ubiquitin | 50, 25, 12.5, 6.25, 3.13, 1.56 | $7.15e^{-6}$ |
| Heimdall UEV-Vps23 $UEV_{1–131}$ | *H. sapiens* ubiquitin | 50, 25, 12.5, 6.25, 3.13, 1.56 | $6.09e^{-5}$ |
| Hel UEV-Vps23 $UEV_{1–132}$ | Hel ubiquitin | 50, 25, 12.5, 6.25, 3.13, 1.56 | $2.97e^{-6}$ |
| Hel UEV-Vps23 $UEV_{1–132}$ | *H. sapiens* ubiquitin | 50, 25, 12.5, 6.25, 3.13, 1.56 | $3.07e^{-5}$ |

The UEV sequences are from *Saccharomyces cerevisiae* Vps23 (NCBI accession number NP_013061.1), Odinarchaeota UEV-Vps23 (GCA_001940665_1___MDVT01000007.1_1083), Wukongarchaeia (GCA_016840725_1___JAEORU010000005.1_18), Heimdallarchaeia UEV-Vps28 (GCA_016839875_1___JAEOTK010000137.1_34), Heimdallarchaeia UEV-Vps28 (GCA_016839875_1___JAEOTK010000223.1_89), and Helarchaeia UEV-Vps23 (GCA_005191425_1___SUPR01000004.1_60). The ubiquitin sequences are from Odinarchaeia (GCA_001940665_1___MDVT01000007.1_1080), Wukongarchaeia (GCA_016840725_1___JAEORU010000005.1_21), Heimdallarchaeia (GCA_016839875_1___JAEOTK010000137.1_32), and Helarchaeia (GCA_005191425_1___SUPR01000004.1_63).

**Table S2.**

**Surface plasmon resonance analysis of interaction between UEV-Vps23 UEV and RING E3 in Helarchaeia.**

| Immobilized on CM5 chip | Injection | Concentration (μM) | Ka 1 (1/Ms) | $K_D$ 1 (1/s) | Ka 2 (1/s) | $K_D$ 2 (1/s) | $K_D$ (M) |
|---|---|---|---|---|---|---|---|
| Hel UEV-Vps23 $UEV_{1–132}$ | Hel RING E3 ($\varDelta$N-loop) | 14, 7, 3.5, 1.75, 0.9 | $2.61e^{+1}$ | $6.16e^{-1}$ | $1.37e^{-2}$ | $7.26e^{-3}$ | $8.14e^{-3}$ |
| Hel Vps23 $UEV_{1–132}$ | Hel RING E3 | 14, 7, 3.5, 1.75, 0.9 | $8.62e^{+1}$ | $5.32e^{-1}$ | $7.68e^{-3}$ | $3.33e^{-4}$ | $2.57e^{-4}$ |

The RING E3 is from Helarchaeia (GCA_005191425_1___SUPR01000004.1_62).

**Table S3.**

**Summary of the eukaryotic and TACK archaeal proteins used in this study.**

| Species | Protein | UniProt accession number |
|---|---|---|
| *Sulfolobus acidocaldarius* | CdvA | Q4J923 |
| *Sulfurisphaera tokodaii* | CdvA | Q972B5 |
| *Stygiolobus azoricus* | CdvA | A0A650CNE5 |
| *Aeropyrum pernix* | CdvA | Q9YFV5 |
| *Fervidicoccus fontis* | CdvA | I0A073 |
| *Ignicoccus islandicus* | CdvA | A0A0U3F871 |
| *Pyrolobus fumarii* | CdvA | G0EE18 |
| *Pyrodictium occultum* | CdvA | A0A0V8RVZ4 |
| *Saccharolobus solfataricus* | CdvA | Q97ZJ5 |
| *Metallosphaera yellowstonensis* | CdvA | H2C8A9 |
| *Acidianus hospitalis* | CdvA | F4B4A8 |
| *Homo sapiens* | Tsg101 | Q99816 |
| *Mus musculus* | Tsg101 | Q61187 |
| *Xenopus tropicalis* | Tsg101 | A0A803JBS5 |
| *Drosophila rhopaloa* | Tsg101 | A0A6P4EG22 |
| *Saccharomyces cerevisiae* | Vps23 | P25604 |
| *Tetrahymena thermophila* | Vps23 | I7MGI0 |
| *Cyanidioschyzon merolae* | Vps23 | M1V8C4 |
| *Chlamydomonas reinhardtii* | Vps23 | A0A2K3DLC5 |
| *Arabidopsis thaliana* | Vps23 | Q9LHG8 |

### Table S4.

### Summary of the Asgard proteins used in this study.

| Protein name | Protein | Species | GCA_name |
|---|---|---|---|
| Helarchaeota_E3 | RING E3 | Helarchaeia | GCA_005191425_1___SUPR01000004.1_62 |
| As_047-p_00441 | RING E3 | Helarchaeia | GCA_005191415_1___SUPS01000012.1_14 |
| As_098-p_02178 | RING E3 | Helarchaeia | GCA_016840125_1___JAEOSY010000201.1_15 |
| As_080-p_02157 | RING E3 | Hermodarchaeia | GCA_016840585_1___JAEOSB010000082.1_4 |
| As_168-p_00310 | RING E3 | Hermodarchaeia | GCA_016550415_1___JABXGU010000063.1_7 |
| As_171-p_01927 | RING E3 | Hermodarchaeia | GCA_016550385_1___JABXGT010000343.1_9 |
| As_097-p_01271 | RING E3 | Hermodarchaeia | GCA_016840135_1___JAEOSX010000144.1_4 |
| As_006-p_01386 | RING E3 | Odinarchaeia | GCA_001940665_1___MDVT01000007.1_1081 |
| As_129-p_01938 | RING E3 | Baldrarchaeia | GCA_016840485_1___JAEOSG010000072.1_7 |
| As_001-p_01350 | Ubiquitin | Heimdallarchaeia | GCA_001940755_1___MEHH01000092.1_2 |
| As_097-p_01272 | Ubiquitin | Hermodarchaeia | GCA_016840135_1___JAEOSX010000144.1_5 |
| As_106-p_02716 | Ubiquitin | Gerdarchaeia | GCA_011366285_1___VIKK01000184.1_10 |
| Helarchaeota_UEV-Vps23 | UEV-Vps23 | Helarchaeia | GCA_005191425_1___SUPR01000004.1_60 |
| TEKIR_9-p_00239 | UEV-Vps23 | Helarchaeia | GCA_004524165_1___SDMZ01000007.1_10 |
| TEKIR_23-p_00034 | UEV-Vps23 | Helarchaeia | GCA_004524205_1___SDNC01000002.1_12 |
| Bin_186-p_02424 | UEV-Vps23 | Helarchaeia | GCA_014729825_1___WJJF01000359.1_1 |
| As_173-p_00118 | UEV-Vps23 | Hermodarchaeia | GCA_016550395_1___JABXGY010000016.1_6 |
| As_171-p_01925 | UEV-Vps23 | Hermodarchaeia | GCA_016550385_1___JABXGT010000343.1_7 |
| As_097-p_01269 | UEV-Vps23 | Hermodarchaeia | GCA_016840135_1___JAEOSX010000144.1_2 |
| As_101-p_00205 | UEV-Vps23 | Hermodarchaeia | GCA_016840065_1___JAEOTB010000010.1_42 |
| LGG330-p_02111 | UEV-Vps23 | Hermodarchaeia | LMSG_G000000617.1___contig_774584_13 |
| SQRJ82-p_02468 | UEV-Vps23 | Jordarchaeia | LMSG_G000000626.1___contig_468509_5 |
| GB128-p_02456 | UEV-Vps23 | Jordarchaeia | LMSG_G000000625.1___contig_3815327_57 |
| LC30-p_00663 | UEV-Vps23 | LC30 | GCA_019058495_1___JAHPYV010000016.1_3 |
| LC20-p_00581 | UEV-Vps23 | LC30 | GCA_019058575_1___JAHPYU010000013.1_9 |
| As_131-p_00043 | UEV-Vps23 | Hodarchaeia | GCA_016839295_1___JAEOUP010000001.1_43 |
| As_051-p_02314 | UEV-Vps23 | Hodarchaeia | GCA_011364965_1___RDOB01000007.1_84 |
| As_082-p_00030 | UEV-Vps23 | Wukongarchaeia | GCA_016840505_1___JAEOSE010000002.1_16 |
| As_075-p_00124 | UEV-Vps23 | Wukongarchaeia | GCA_016840725_1___JAEORU010000005.1_21 |
| As_085-p_00172 | UEV-Vps23 | Wukongarchaeia | GCA_016840425_1___JAEOSI010000007.1_21 |
| As_053-p_01062 | UEV-Vps23 | Gerdarchaeia | GCA_011364945_1___RDOD01000006.1_80 |
| As_134-p_01460 | UEV-Vps23 | Gerdarchaeia | GCA_013166835_1___JABLTI010000046.1_19 |
| As_183-p_00946 | UEV-Vps23 | Borrarchaeia | GCA_016840315_1___JAEOSO010000051.1_6 |
| As_178-p_01875 | UEV-Vps23 | Borrarchaeia | GCA_016840645_1___JAEORY010000106.1_6 |
| As_181-p_00830 | UEV-Vps23 | Borrarchaeia | GCA_016840515_1___JAEOSF010000008.1_67 |
| As_129-p_01940 | UEV-Vps23 | Baldrarchaeia | GCA_016840485_1___JAEOSG010000072.1_9 |
| As_130-p_02275 | UEV-Vps23 | Baldrarchaeia | GCA_016840465_1___JAEOSH010000070.1_9 |

| | | | |
|---|---|---|---|
| As_006-p_01388 | UEV-Vps23 | Odinarchaeia | GCA_001940665_1___MDVT01000007.1_1083 |
| As_143-p_00986 | UEV-Vps23 | Odinarchaeia | GCA_016839265_1___JAEOUO010000001.1_1003 |
| As_047-p_00889 | UEV-Vps23 | Helarchaeia | GCA_005191415_1___SUPS01000031.1_8 |
| As_098-p_02180 | UEV-Vps23 | Helarchaeia | GCA_016840125_1___JAEOSY010000201.1_17 |
| LC20_-p_00581 | UEV-Vps23 | Freyarchaeota | GCA_019058575_1___JAHPYU010000013.1_9 |
| LC30_-p_00663 | UEV-Vps23 | Freyarchaeota | GCA_019058495_1___JAHPYV010000016.1_3 |
| As_001-p_01333 | UEV-Vps23 | Heimdallarchaeota | GCA_001940755_1___MEHH01000089.1_12 |
| As_139-p_01042 | UEV-Vps23 | Heimdallarchaeota | GCA_013166775_1___JABLTJ010000016.1_8 |
| As_017-p_02597 | UEV-Vps23 | Kariarchaeota | GCA_016839545_1___JAEOUB010000567.1_6 |
| S012_26_esom-p_00393 | UEV-Vps23 | Kariarchaeota | GCA_015523565_1___WAKA01000032.1_6 |
| H2.bin.2_-p_03262 | UEV-Vps23 | Kariarchaeota | GCA_021399405_1___JAAFKL010000319.1_6 |
